# High-Speed Multiplexed DNA-PAINT Imaging of Nuclear Organization using an Expanded Sequence Repertoire

**DOI:** 10.1101/2024.12.21.629871

**Authors:** Abhinav Banerjee, Micky Anand, Mansi Srivastava, Vedanth Shree Vidwath, Mahipal Ganji

**Author notes:** Equal contribution (Abhinav Banerjee, Micky Anand).

## Abstract

DNA-Points Accumulation for Imaging in Nanoscale Topography (DNA-PAINT) enables multiplexed super-resolution imaging of biological samples. We expand the repertoire of speed-optimized DNA sequences for DNA-PAINT imaging to drive visualization of as many as twelve targets in a sequential manner with molecular resolution. By implementing Exchange-PAINT protocol, we demonstrate 12-plex super-resolved imaging of docking strand patterned DNA origami nanostructures within four hours with a localization precision of 3 to 5 nm. Using these sequences, we demonstrate 9-plex super-resolution imaging of diverse nuclear targets within four hours. Further, we present a comprehensive analysis pipeline to quantify nanoscale chromatin in single cells. The combination of multiplexed imaging and analysis pipeline enabled us to reveal the loss of chromatin contacts with nuclear speckles upon global transcription inhibition. This work highlights the versatility of our approach to simultaneously image multiple targets at accelerated speeds while maintaining precise spatial localization for each target, enabling in depth mapping of the nuclear landscape. These speed-optimized imager sequences for high multiplexed super-resolution imaging will drive its further adoption for diverse cellular imaging applications.

## Introduction

Recent years have witnessed tremendous development in a variety of super-resolution imaging techniques, which enable us to gain greater insights into the spatial distribution of biomolecules at nanometer resolution in intact cells^1-4^. DNA-PAINT, a variant of Single-Molecule Localization Microscopy, achieves diffraction unlimited resolution through stochastic, transient hybridization of fluorophore-tagged short complementary single-stranded DNA molecules.^2-5^ Additionally, DNA-PAINT enables multiplexed imaging by encoding the target identity into DNA docking strands. However, the acquisition times are enormous, requiring up to several hours for each imaging round, because of the rather slow binding kinetics of DNA sequences used during the infancy of DNA-PAINT^6,7^. Recent studies have improved imaging speed by cleverly designing imager strands with non-complementary nucleotides arranged in repetitive blocks forming concatenated sequences, thereby increasing the frequency of imager-docking strand hybridization.^8^

Currently, only six such speed-optimized sequences are available, limiting multiplexing to six targets. Although recent approaches using secondary^9^ or adapter DNA probes^10^ allow near-unlimited multiplexing capabilities, they require complex strand design to avoid non-orthogonality along with tedious and often inefficient hybridization and dissociation schemes.^11^ Thus, expanding the sequence space to enable faster imaging with greater multiplexing-ability remains a key challenge.

In this manuscript, we report six additional imager sequences hence increasing the multiplexing capabilities of speed optimized DNA-PAINT to 12 target imaging in a single experiment. The new sequences that display longer binding times show additional benefits of higher achievable resolution and faster target sampling. Using these sequences, we performed 9-plex nuclear imaging to reveal overall chromatin organization in single cells. In addition to mapping the spatial chromatin landscape, we observe a profound change in contacts between different chromatin marks and nuclear speckles upon transcription inhibition. Our study emphasizes the applicability of speed-optimized sequences for obtaining multiplexed spatial organization of biomolecules in single cells.

## Results and Discussion

### Design principles for generating speed optimized DNA-PAINT sequences

To outline sequences that could be used for speed-optimized DNA-PAINT, we systematically explored all combinations of orthogonal imager-docking strand pairs in the entire sequence space that support faster imaging. Two fundamental design rules that were followed in this search are that (1) the imager strand must not comprise nucleotides that are complementary, and (2) they must form core units that can be concatenated to form overlapped docking strands^8^ (Supplementary Figure 1a, b). This yielded two di-, four tri-, and six tetra-nucleotide blocks (Figure 1a, Supplementary Figure 1c). We reserved cytosine (C) containing blocks for docking strands, and guanine (G) containing blocks for imagers. The presence of non-complementary nucleotides in these sequences preclude any secondary structures, ensuring that the DNA strands remain in an open configuration. This conformation facilitates rapid and efficient binding to the complementary strand. Concatenation of such blocks in the docking strand further improves binding rates by increasing the number of potential binding sites for the imager.^12^ This design strategy yielded as many as 12 sequences of which six sequences were reported earlier^8^ (Figure 1b). Increasing the block length to five nucleotides compromises orthogonality and was hence excluded from further consideration (Supplementary Figure 1d).

**Figure 1.**
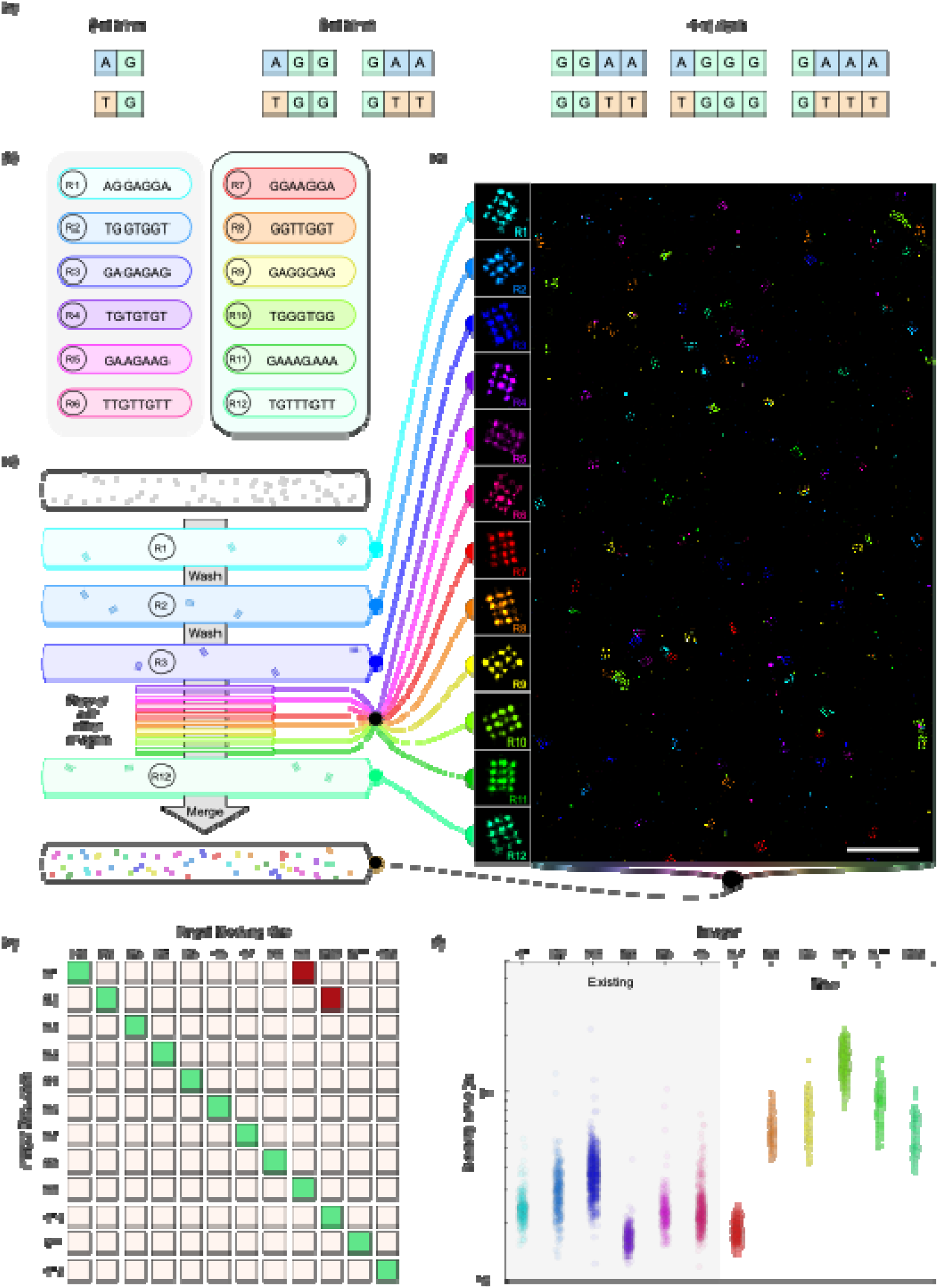
Full repertoire of imager strands for high-speed DNA-PAINT and their binding kinetics. (a) All possible building blocks for constructing imager strands. (b) The sequences of all 12 imager strands. R1 to R6 imagers have been published previously^8^. (c) Scheme for 12-color multiplexed imaging using Exchange-PAINT. (d) Merged 12-color DNA-PAINT data of DNA origami nanostructures. 10000 frames were captured for each imaging channel in 1000 seconds. (e) Chart showing orthogonality and cross-binding of imagers. Red squares depict unexpected cross-binding (non-target origamis showing signal greater than 10% of the target origami). Green squares depict specific binding. (f) Scatter plot showing average of individual binding times per binding site of each docking strand. In (b) and (f), ‘existing’ refers to imagers reported earlier.^8^ (Scale: (d) 500nm)

We tested the multiplexing capability of all 12 speed-optimized sequences utilizing unique DNA origami nanostructures carrying docking strand extensions for each of the imager sequences. For visual identification and testing the resolution, we designed origami nanostructures carrying docking strands in a 4×3 grid pattern where each strand is separated from its neighbor by 20 nm. We immobilized these in a single microfluidic channel and imaged them in 12 rounds using Exchange-PAINT modality^5^ (Figure 1c). Each species was imaged for 1000 seconds, with washes entailing 300 seconds between each round, taking less than 4.5 hours for 12-plex imaging. As expected, all 12 species of DNA origami nanostructures were clearly visible in the merged data with a localization precision lower that 5 nm (Figure 1d and Supplementary Figure 2). Ten of the twelve imagers showed clear specific binding on their target docking strands in an orthogonal fashion. Two imagers, R1 and R2, displayed unexpected non-specific binding, with R1 binding to R9 origami structures and R2 binding to R10 origami structures (Figure 1e, Supplementary Figure 3 and Supplementary Table 1). This emphasizes the tolerance for certain DNA mismatches to form transient duplexes with sufficient stability allowing for detection of a binding event. This unexpected non-specific binding further highlights the limitations of secondary probe-based multiplexing approaches where longer probes are more prone to unpredictable non-orthogonality. Because of this cross-binding, only one imager from each of these pairs can be used for multiplexed imaging to maintain orthogonality. Together, Exchange-PAINT imaging of “12-color” data demonstrates that we can achieve multiplexing capabilities of as many as ten targets at high-speed and high-resolution.

We next performed kinetic analysis of all twelve imager-docking strand pairs by utilizing DNA nanostructures with patterns carrying a unique docking strand in the center (Supplementary Figure 4a). We performed two-color DNA-PAINT imaging to obtain the kinetics of the docking strand first, followed by the pattern of the nanostructure (Supplementary Figure 4b). Our analysis shows that these imagers display binding times ranging from a few hundred milliseconds to over a second (Figure 1f and Supplementary Figure 4c). This versatility in range of binding times enables us to utilize imager strands based on the required properties for a given imaging application. For instance, imagers with wide range of kinetics can enhance the multiplexing capabilities of DNA-PAINT via kinetic barcoding^13,14^. The speed of DNA-PAINT imaging is determined by the frequency of hybridization events between the imager and its corresponding docking strand, a parameter that we systematically analyzed. We observed that the average time between two consecutive binding events (dark time) of these imagers ranged between 10^2^ to 10^3^ seconds on a single docking strand repeat at 1 nM imager concentration which is in agreement with the previous report (Supplementary Figure 4c). The dark times can be further tuned by modulating the length of concatenated docking strands. Short dark times allow for a higher frequency of binding events at lower imager concentrations reducing fluorophore background while concurrently improving sampling and the resolution of the image.

### Long binding imagers have distinct advantages

Five of the six (R8 – R12) newly proposed imagers show longer binding times as compared to the other speed-optimized sequences (Figure 1f and Supplementary Figure 4c). These imagers have a few notable advantages over the shorter binding imagers. Firstly, with long binding imagers, one can achieve better imaging resolution (localization precision) owing to greater number of frames with binding events spanning the entire frame as compared to those showing shorter binding times (Figure 2a). The full frame events have better signal-to-noise ratio compared to partial frame events and thus improving the localization precision. We compared the ability of two imagers with different binding times: R1 with an average time of 282 milliseconds, and R8 with an average time of 678 milliseconds to resolve the 12 docking strand extensions (4×3 grid) separated by 20 nanometers arranged on a DNA origami nanostructure. At a low laser power density, imaging with R1 does not resolve the individual docking strands. On the other hand, the R8 imager resolves the 4×3 grid with α= 5.71 nm, the standard deviation of the fitted Gaussian distribution to the localization signal, representing localization precision. At a high laser power density, R1 is able to resolve the grids with α= 5.16 nm, whereas R8 resolves them with a considerably better precision of α = 3.79 nm (Figure 2b). This showcases the advantages of longer binding imagers in achieving better resolution under both lower and higher laser power densities.

**Figure 2.**
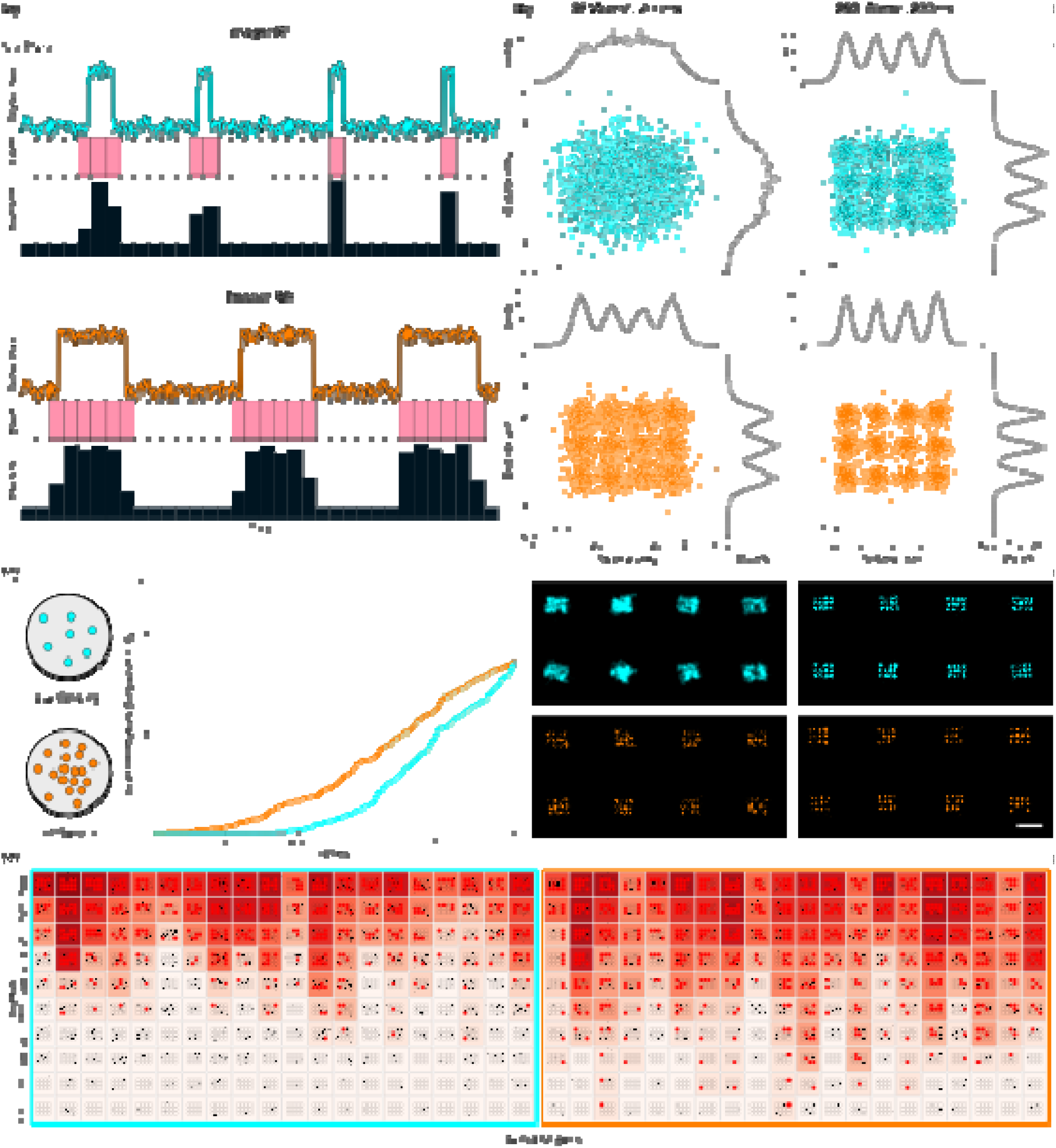
Longer binding imagers provide better localization precision and target sampling. (a) Schematic binding time traces for short (top) and long (bottom) binding imagers. (b) Top: Average of patterned DNA origami nanostructure imaged with R1 (cyan) and R8 (orange) at 37 W/cm^2^ (left) and 250 W/cm^2^ (right). Bottom: Representative origami nanostructures imaged with R1 (cyan) and R8 (orange) at 37 W/cm^2^ (left) and 250 W/cm^2^ (right). (c) Fraction of targets sampled over time for R1 (cyan) and R8 (orange). (d) Time evolution heat maps of example origamis, left is R1 and right is R8. Solid red dots indicate a complete sampling with more than 20 localizations. (Scale: (b) 100nm)

The long binding imagers provide additional flexibility by allowing imaging at lower laser power densities and slower frame rates (longer exposure). This allows the camera to collect more photons per binding event resulting in a localization precision comparable to that obtained with a short binding imager at ∼8-fold higher laser power density. This is crucial as higher laser powers often lead to laser induced damage and loss of docking strands over time.^15^ Additionally, long binding imagers would generate a greater number of localizations at the same imaging frame rate upon each binding event. This would improve target sampling and thus require shorter acquisition times to adequately sample the targets at similar depths. We compared the target sampling on DNA origami nanostructures carrying 12 docking strands in a4×3 grid. Considering a threshold of 20 localizations to sample a given target site, we observe that R8 shows faster sampling as compared to R1 while having comparable dark times (Figure 2c). Individual origami heatmaps confirm this trend across many origami structures (Figure 2d) making longer binding imagers faster at target sampling.

### New sequences are optimal for cellular imaging

To establish the ability of new sequences to image cellular targets, we tested their behavior by imaging the nuclear lamina (Lamin B1) and found that all the imagers showed the expected signals, except R3, which displayed additional non-specific nucleolar localizations (Supplementary Figure 5, 6 and 7). The data reveals that accelerated multiplexed imaging of cellular targets with an Exchange-PAINT based approach is readily achievable with the expanded sequence repertoire.

The true power of multiplexing can be exploited only when many cellular targets are visualized to map their spatial distribution within the cell. Earlier strategies to build a super-resolved map of many proteins were performed using non-optimized DNA-PAINT sequences, showcasing slow on-rates, thus requiring massive acquisition times^16,17^. With the development of new speed-optimized sequences, we can now push the boundaries of multiplexed cellular imaging to more than six targets in shorter time scales.

### High-Speed Multiplexed Imaging of 9 Nuclear Targets with Exchange-PAINT

With our ability to perform direct multiplexed imaging of cellular targets, we set out to build a spatial map of the chromatin landscape in a mammalian nucleus. A significant challenge in conventional immunofluorescence-based imaging of nuclear targets is the diffuse signal obtained for the target because of their distribution across the entire nuclear volume. We strategized to exploit these sequences to delineate the nanoscale spatial distribution of various nuclear features. The nuclear environment poses an unavoidable challenge to DNA-PAINT due to the abundance of chromatin polymer which can often lead to spurious off-target binding events resulting in a homogenous artefact in the data. Due to the propensity of R3 imager to generate off-target nucleolar localizations, it was omitted from the multiplexing imager pools. We tested all the imagers for their propensity to generate off-target nuclear localizations. In absence of a corresponding primary antibody, the docking strand-conjugated nanobody alone controls showed sparse nuclear localizations. In contrast, the presence of primary antibodies showed prominent nuclear localizations. (Supplementary Figure 8, 9 and 10). These data support the applicability of imager sequences for multiplexed imaging in the challenging environment of the nucleus.

We targeted nine different nuclear marks encompassing features known to delineate actively transcribed euchromatin (RNA Pol II Serine 2 Phosphorylation (RNAPolII-S2P), RNA Pol II Serine 5 Phosphorylation (RNAPolII-S5P), splicing factor SC35, H3K4me3 and H3K37ac); heterochromatin (H3K27me3, H3K9me3, Lamin B1); and CTCF, a transcription factor involved in dictating genome architecture. For achieving multiplexed imaging with the same species primary antibodies, we preincubated them with secondary nanobodies, each conjugated to a unique DNA docking strand (Figure 3a).^9,10^ This strategy avoids cross-reactivity by encoding a unique DNA conjugated nanobody to each primary antibody enabling us to visualize multiple nuclear targets using DNA-PAINT. We were able to capture differential spatial distribution of all targets relative to each other showcasing the complexity of nuclear organization in a single cell captured through multiplexed imaging (Figure 3b, Supplementary Figure 11 to 16).

**Figure 3.**
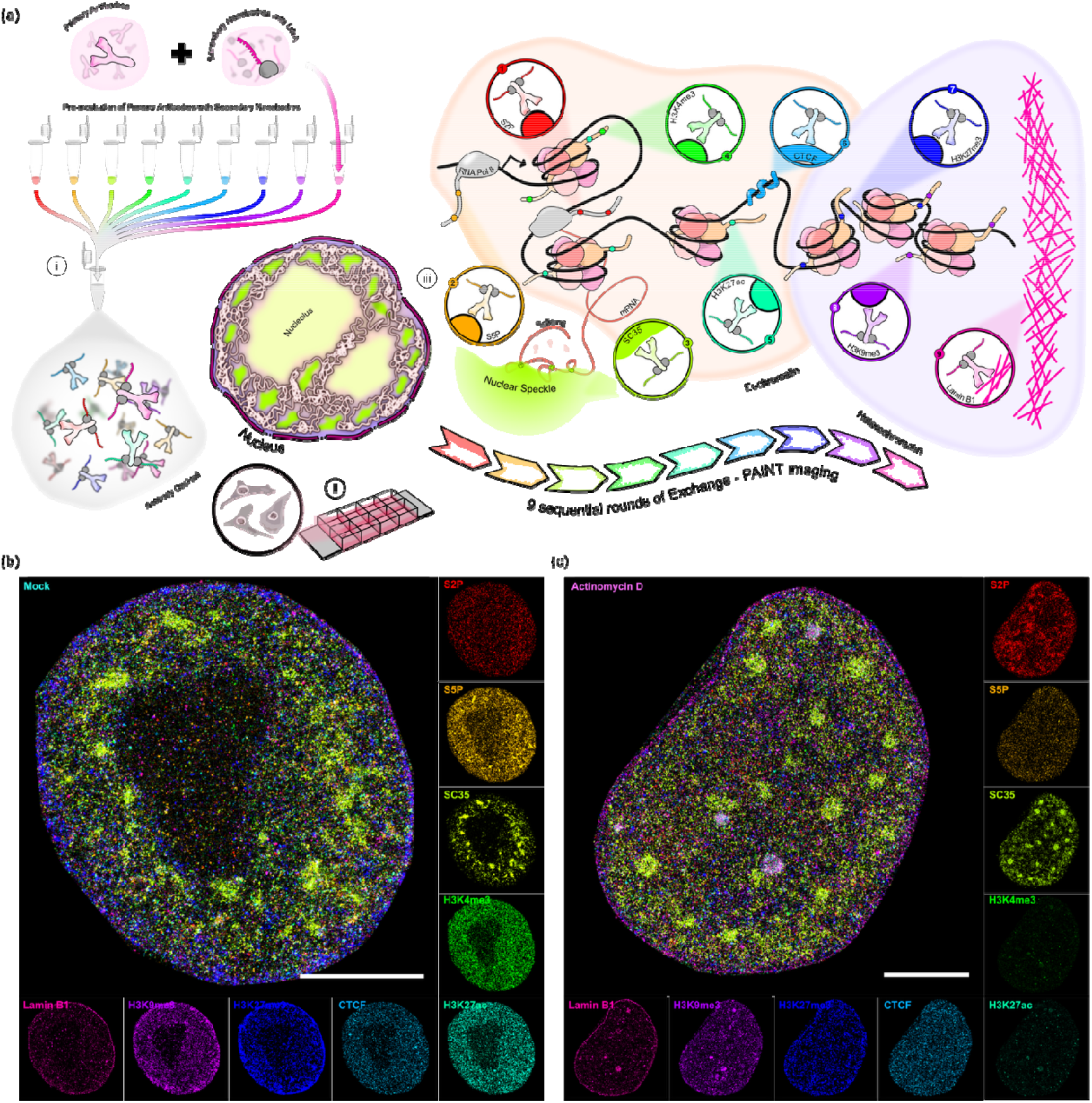
9–plex imaging of the nucleus. (a) Experimental scheme. (i) Preincubation of primary antibodies with DNA conjugated secondary nanobodies. (ii) Cells grown in glass bottom 8-well dishes. (iii) Nine nuclear marks imaged in sequential imaging rounds; Nine-plex images of single nuclei showing detailed organization of targets in (b) mock (left) and (c) ActD (right) treated cells. (Scale: (b) 5 µm)

To quantitatively interpret our data, we employed a coarse-grained correlation analysis between target pairs (see Methods) comparing the number of localizations in each pixel across different protein pairs. We observe expected associations among targets in the actively transcribed regions in DMSO-treated (mock) cells.^18^ For instance, RNAPolII-S2P demarcating elongating transcriptional machinery and initiation primed RNAPolII-S5P show positive correlation with the nuclear speckles (SC35), highlighting their physical proximity. ^18^ For instance, RNAPpolII-S2P demarcating elongating transcriptional machinery and initiation primed RNAPpolII-S5P show positive correlation with the nuclear speckles (SC35), highlighting their physical proximity. Similarly, heterochromatin marks H3K27me3 and H3K9me3 show expected positive correlation with the nuclear lamina (Lamin B1) and negative correlation with the nuclear speckles and the RNA polymerase marks. (Figure 4a–below diagonal, Supplementary Figure 11 to 16). We also observed expected mutual exclusion of euchromatin and heterochromatin marks.

**Figure 4.**
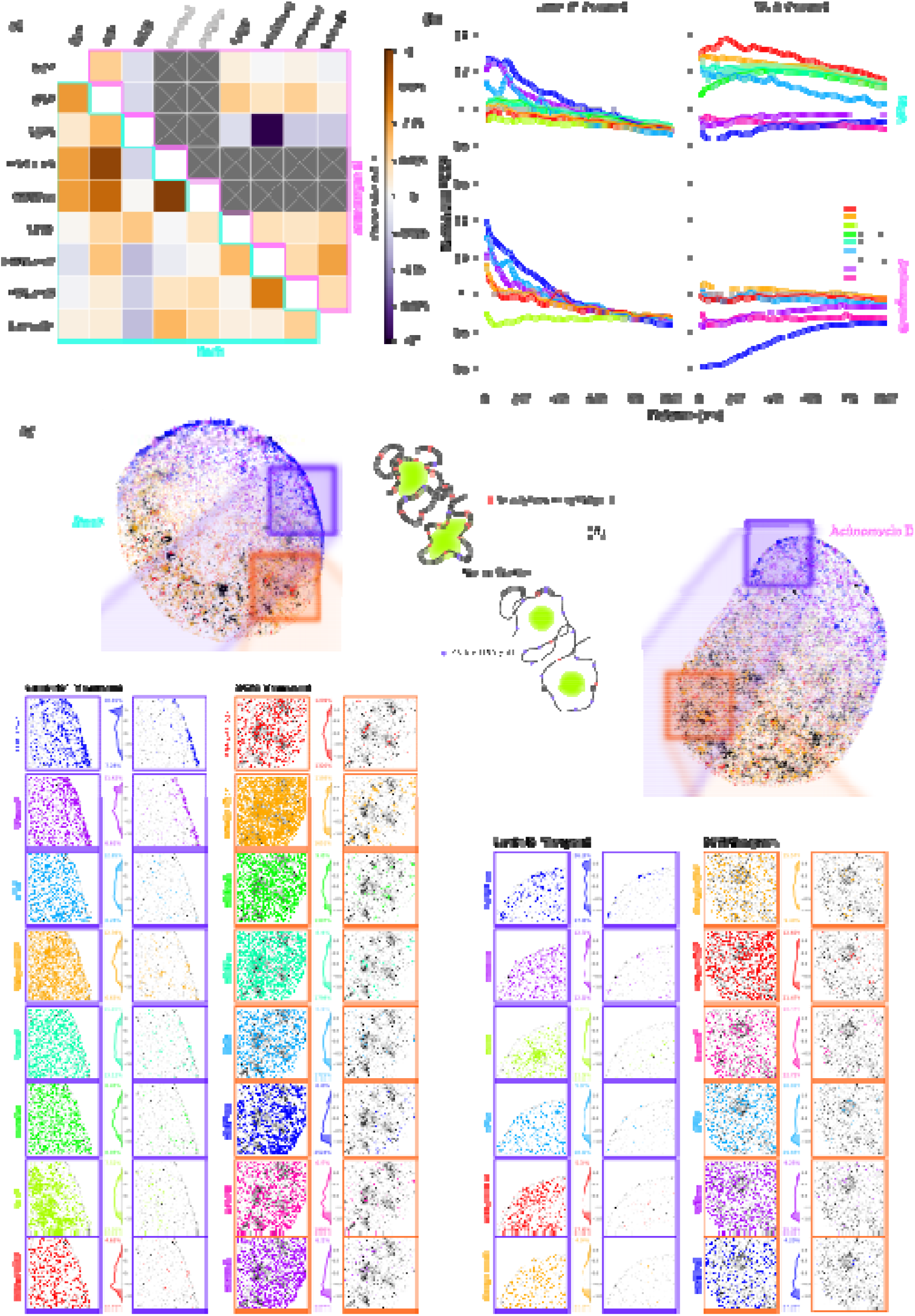
Transcription inhibition perturbs nuclear organization at multiple scales. (a) Global Pearson’s correlation heatmap showing spatial relationships among nine nuclear marks in control (Mock, cyan) and transcription inhibited cells (ActD, magenta) cells (b) PCCF analysis of Mock (top) and ActD (bottom) treated cells from the viewpoint of Lamin B1 (left) and SC35 (right). Viewpoint analyses of spatial correlations in (c) Mock and (d) ActD treated cells. Full cells: Localizations with high positive correlation (ρ > 0.7) of H3K27me3 (Blue) and H3K9me3 (Violet) with Lamin B1 (Black) in the Lamin viewpoint sections, and S2P (Red) and S5P (Golden-Yellow) with SC35 (Black) in the SC35 viewpoint, respectively. Zoom Ins: Left column – all localization for the respective target (in color) and localizations for Lamin B1 or SC35 (in black), respectively. Centre column – KDE plots of the spatial correlations for all localization of the target with the reference (Lamin B1 and SC35 respectively) in left column. Shaded portions show high positive and negative correlations (*ρ* > 0.7 and *ρ* < −0.7). Percentage of points with high spatial correlation is outlined. Right column – spatial distribution of localizations with high positive correlation (ρ > 0.7) with the reference, respectively. Looped models represent likely chromatin re-distribution around nuclear speckles upon transcription inhibition.

### Transcription inhibition leads to large scale restructuring of the chromatin landscape

Having established the ability to spatially distinguish closely spaced nuclear targets, we were interested in visualizing the redistribution of these nuclear marks under perturbation of nuclear processes. Transcription is one such process that can alter the local chromatin organization. We tested if our multiplexing approach could detect such rearrangements under transcriptional inhibition. We treated cells with Actinomycin D (ActD), a known drug that stably intercalates in the genomic DNA and inhibits transcription.^19^ We detected profound nanoscale changes in the chromatin organization upon ActD treatment. Notably, nuclear speckles appeared more circular compared to the amorphous structures observed under mock conditions. The amorphous speckles arise in the untreated cells via chromatin tethering to nuclear speckles because of co-transcriptional splicing^20^. However, transcription inhibition by ActD halts new RNA synthesis, disrupting the chromatin tethering, hence the nuclear speckles morphology appears as circular (Figure 3c, Supplementary Figures 17 to 23).

Interestingly, inhibition of transcription by ActD leads to reduction in overall levels of H3K27ac and H3K4me3^21,22^ among all the targets imaged. We confirmed this result by individual targets imaging after ActD treatment (Supplementary Figures 24 to 26). We therefore omitted H3K27ac and H3K4me3 for further correlation analysis for ActD treated cells (Fig 4a-top diagonal, Supplementary Figures 17 to 23). While the nuclear speckles positively correlated with euchromatin marks in untreated cells, inhibition of transcription lead to negative correlation between speckles and all chromatin modifications (Figure 4a-top diagonal, Supplementary Figures 17 to 23). For instance, heterochromatin marks displayed increased overall positive correlation with the nuclear lamina. These results establish the robustness of the coarse-grained analysis in capturing the chromatin redistributions in a single cell.

Because nucleus is a highly dense and inhomogeneous compartment containing diverse nuclear bodies, global correlation analysis produces averaged correlations that obscure specific local interactions. To overcome this limitation, we performed spatial correlation analysis between each pair of targets. For this, we computed the pairwise cross-correlation function (PCCF) between target pairs. The PCCF is a measure of the probability of finding a target at a certain distance with another target when compared to a random distribution. A PCCF value of one signifies a random distribution of the target with respect to the other target used as the viewpoint. A value higher than one at a given spatial distance signifies the spatial correlation or interaction between the two targets. A value lower than one indicates a negative spatial correlation or mutual exclusion between the targets. We observed a clear positive correlation between all four euchromatin markers and SC35. Concurrently, a negative correlation was observed for heterochromatin markers and nuclear lamina from SC35 viewpoint (Figure 4b-SC35 viewpoint). On the other hand, from the lamina point of view, heterochromatin marks displayed a positive correlation, whereas both speckles and euchromatin showed diminished interactions with it (Figure 4b-Lamin B1 viewpoint).

To further understand the spatial organization of the targets in the nucleus, we calculated the Spearman’s correlations between two targets encompassed by concentric donuts centered over a given localization^23^. This measurement, also called DoC (Degree of Colocalization), provides a suitable measure of spatial correlation that can be attributed to each localization^23^. Such spatial correlation maps enable us to discern regions where interactions are significantly higher and correlate better with the target of interest. We selected two targets with defined physical features for this purpose: 1) nuclear lamina and 2) nuclear speckles, representing heterochromatin and euchromatin region, respectively.

This analyses once again embodies the expected correlation amongst heterochromatin and euchromatin regions. Heterochromatin marks show a clear mutual interaction. For instance, H3K27me3 and H3K9me3 showed strong positive correlation (>0.7) with the nuclear lamina. Notably, SC35 displayed a higher fraction of negative correlation with the RNA polymerase markers (Figure 4c–SC35 Viewpoint). We note that the DoC analysis contradicts with PCCF analysis where SC35 and euchromatin showed positive correlations. This apparent contradiction likely arises from the peripheral arrangement of active transcription and chromatin around nuclear speckles.^24^ This arrangement leads to decreased localizations of SC35 in the concentric donuts around the periphery of speckles with a corresponding increase in localizations of the active transcription and chromatin marks, resulting in the negative Spearman’s correlation or DoC between them.

## Conclusions

We present an expanded set of 12 speed-optimized DNA-PAINT sequences. Using designer DNA origami structures, we characterized their orthogonality and sampling rates for obtaining high-resolution images. The sequences display accelerated hybridization rates reducing acquisition times from hours to a few minutes at comparatively lower imager concentrations. Using these new imager sequences, we demonstrate 9-plex nuclear imaging in just under 4 hours using Exchange-PAINT with a localization precision as good as 8 nm. In untreated cells, we observed expected positive spatial correlation between euchromatin marks (H3K27ac and H3K4me3), active transcription marks, and nuclear speckles (SC35). As expected, the euchromatin marks showed negative correlation with the heterochromatin marks and the nuclear lamina. Inhibition of transcription using ActD led to a global restructuring of the chromatin landscape. We observed a loss of correlation between SC35 with all the chromatin marks consistent with the loss of RNA mediated tethering between chromatin and speckles. Additionally, heterochromatin interactions with the nuclear lamina become stronger with the concurrent reduction in the levels of euchromatin marks. To capture local chromatin interactions, we performed PCCF analysis which revealed positive spatial correlation of euchromatin with nuclear speckles, and heterochromatin with nuclear lamina.

This expanded pool of speed-optimized imager-docking strand sequences will unlock the full potential of DNA-PAINT imaging’s multiplexing capabilities. Additionally, the adapter-strand-based “unlimited” multiplexing can be done in batches of ten instead of six targets per labeling round which would drastically reduce the time required for multiplexing. The increased repertoire of speed-optimized imager-docking strand sequences will also facilitate multi-color sub-nanometer resolution imaging with RESI which requires multiple rounds of imaging for each target identification.^25^ In addition, by combining with orthogonality of left-handed DNA imagers, multiplexing can be extended to as many as 18 targets.^26^ Simultaneous imaging of histone modifications, transcriptional machinery, and nuclear bodies at high resolution, signify the application of new sequences in the field of spatial proteomics and interactome building, in addition to other allied fields including biomedical sciences.

## Acknowledgements

We acknowledge Vinay Bhardwaj for helping with purifying anti-rabbit IgG secondary nanobody. A.B. and M.A. acknowledge the support from the Prime Minister’s Research Fellowship (PMRF), Ministry of Education, Government of India. We greatly acknowledge Sarit Agasti for allowing us to use their laboratory facilities and S. Kalita for help with HPLC purification of DNA imager strands.

