## Supplementary material for "High-Speed Multiplexed DNA-PAINT Imaging of Nuclear Organization using an Expanded Sequence Repertoire": Methods and materials

Buffers

10× Folding Buffer: 500 mM Tris-Cl pH 8.0 (Tris-Base, Sigma #77861; HCl, Fisher Scientific #29507), 125 mM MgCl_2_ (EMPLURA #105833) and 2 mM EDTA (SRL #35888).

Buffer A: 10 mM Tris-Cl pH 8.0 and 100 mM NaCl (SRL #3205)

Buffer A+: 10 mM Tris-Cl pH 8.0, 100 mM NaCl and 0.05% v/v Tween 20 (Sigma #P9416-100ML)

Buffer I+: 50 mM Tris-Cl, pH 8.0, 10 mM MgCl_2_, 0.2 mM EDTA, 0.05% v/v Tween 20.

Lysis Buffer: 50 mM Tris-Cl pH 7.5, 300 mM NaCl, 10 mM Imidazole (SRL #61510) and 5 mM DTT (SRL #84834).

Elution Buffer: 50 mM Tris-Cl Ph 7.5, 150 mM NaCl, 500 mM Imidazole and 10% Glycerol (SRL #77453)

Maleimide Labelling Buffer: 100 mM Sodium Phosphate (NaH_2_PO_4_ SRL #59443, Na_2_HPO_4_ Merck #61795105001730) pH 7.5, 150 mM NaCl and 250 mM Sucrose (EMPLURA #1.94953.0521)

Low Salt Buffer: 1× PBS (SRL #78529).

High Salt Buffer: 1× PBS with 1 M NaCl.

PEM Buffer: 80 mm PIPES (SRL #49159), 5 mm EGTA (SRL #62858), 2 mm MgCl_2_, pH 6.8.

Cytoskeleton Extraction Buffer: 0.25% Triton X-100 (Sigma #T8787-250ML), 0.1% Glutaraldehyde (EMS #16020) in PEM, pre-heated to 37 °C.

Cytoskeleton Fixation Buffer: 0.25% Triton X-100, 0.5% Glutaraldehyde in PEM, pre-heated to 37 °C.

PFA Fixation Buffer: 4% Paraformaldehyde (EMS # 15710) in 1× PBS pre-heated to 37 °C.

Quenching Buffer: 0.1 mg/ml of NaBH_4_ (Sigma #213462-25G) in 1× PBS.

Blocking Buffer: 3% w/v BSA (Sigma #A4503-50G), 0.05 mg/ml Salmon Sperm DNA (Invitrogen #15632011) and 0.25% v/v Triton X-100 in 1× PBS.

Antibody Incubation Buffer: 3% w/v BSA, 0.05 mg/ml Salmon Sperm DNA, 0.02% v/v Tween-20 and 1 mM EDTA in 1× PBS.

Buffer C: 1× PBS supplemented with 500 mM NaCl.

Flow Cell Preparation

Slides and coverslips for microscopic imaging of DNA origami nanostructures were prepared as previously described.^1^ Slides (VWR #631-1550) were drilled with five pairs of holes using a diamond head drill bit (Meisinger #801-009-HP) using a drill gun (DigitalCraft #LRUXOR) mounted on a drill stand (Dremel #220-01 Workstation™) under a shallow water bath. Slides were then washed thoroughly using 5% v/v dish washing detergent to remove any trace stains and contaminants. The slides were then thoroughly rinsed with Milli-Q® water followed by 70% ethanol and once again with Milli-Q®. Coverslips (Marienfeld #0107222) were cleaned similarly prior to use. Flow cells were assembled using double sided tape (3M Scotch #136D MDEU) and the open ends were sealed with quick setting epoxy (Araldite Klear). These flow cells were used for all experiments where DNA origami nanostructures were used.

For multiplexed origami imaging, the flow cells were attached with tubing and syringes for ease in buffer exchange.

DNA Origami Nanostructure Folding

Picasso Design^2^ was used to get all the staple sequences (Supplementary Table 1). M13mp18 ssDNA (Bayou Biolabs #P107) was used as the scaffold ssDNA for folding all the DNA origami nanostructures. Oligonucleotide staples containing biotin at 5′-end (Supplementary Table 1) were used to anchor the DNA origami nanostructures onto the surface. A 30 µl of 1× folding buffer folding reactions were set in PCR Tubes (BioRad #TFI0201) with 10 nM of scaffold DNA, 100 nM of biotin staples, 100 nM of blank staples, 1 µM of staples with docking strands. The mix was then heated up to 80 °C and held for five minutes before a stepwise cooling of 0.1 °C every five seconds till 4 °C in a thermocycler (Applied Biosystems™ ProFlex™ PCR System #4484073).

The folded DNA origami nanostructures were purified from free staples using concentrator centrifugal spin filters (Sartorius Vivaspin® 500 #VS0132). The filters were equilibrated at 3000 ×g for 5 minutes with 500 µl of Milli-Q® water followed by 500 µl of 1× folding buffer. Following this, the origami mix was applied on the column along with 470 µl of 1× folding buffer and spun at 3000 ×g till the volume is reduced to about 50 µl. A 450 µl of 1× folding buffer was then added, and the process was repeated three times to remove the unincorporated staples. The origami structures were then recovered and stored at -20 °C for later use.

Imager Fluorophore Conjugation and Purification

Labeling of DNA strands with fluorophore for preparing imager strands were done by following the published protocol.^1^ The imager sequences were purchased from Merck carrying amine modification at their 3′-end (Supplementary Table 1) and dissolved in Milli-Q® water.

Fluorophores were dissolved in DMSO (Sigma #D8418) and stored at -20 °C at the below mentioned concentrations. Cy3B-MonoNHS-Ester (Cytiva #PA63101) was dissolved at 13 mM concentration. Atto647N-MonoNHS-Ester (Sigma #18373-1MG-F) was dissolved at 11.8 mM concentration.

A 15 nanomoles of DNA (15 µl of DNA stored at 1 mM stock) in 1× PBS and 0.1 M NaHCO_3_ (Sigma #S5761) was used for labeling in each reaction. The labeling was done by using five-fold excess of Cy3B-MonoNHS-Ester or Atto647N-MonoNHS-Ester. The reaction was then left overnight on a shaker at 4 °C.

The incubation mix was then purified to separate the free fluorophore and the unconjugated DNA oligonucleotides from the conjugated product using reverse phase (Phenomenex #00B-4442-E0 Clarity® 5 µm Oligo-RP, LC Column 50 × 4.6 mm, Ea) HPLC (Agilent Technologies). The conjugated product was then lyophilized and then dissolved in Milli-Q® and then stored at -20 °C for future use.

Sample Preparation for DNA Origami Nanostructure Imaging

Microfluidic flow cells were created as mentioned previously.^2^ In Brief, the channels were first incubated with 10 to 15 µl of 1 mg/ml Biotinylated Bovine Serum Albumin (Sigma #A8549) diluted in Buffer A. This was followed by a thorough wash of the channel with Buffer A+. A 10 to 15 µl of 0.2 mg/ml of Neutravidin (Sigma #31000) in Buffer A was then introduced and incubated for a minute. Channels were then washed with Buffer A+ thoroughly. Buffer I+ was used for washing the channel prior to its introduction. Origami nanostructures were then diluted appropriately in Buffer I+ and then incubated on the flow cell for 20 minutes. Thorough washing with 600 – 800 µl of Buffer I+ was performed to remove any unbound DNA origami structures.

Flow cells were incubated with Gold Nanoparticles (Sigma #753688-25ML) at a dilution of 1:10 in buffer I+ for 10 minutes and then washed with 600 – 800 µl of Buffer I+ prior to imaging.

Antibody Conjugation

Secondary antibody against-mouse IgG (Jackson #715-005-150) was purchased. The antibodies were buffer exchanged to 1× PBS and concentrated to over 1.5 mg/ml using centrifugal concentrators as mentioned above. Spins were performed at maximum recommended protocol of 12000 × g. This ensures better conjugation between the crosslinker and the antibody. DBCO-PEG4-NHS Ester crosslinker (Jena Biosciences #CLK-A134-10) stock solution was stored at a concentration of 15.39 mM. A 25-fold excess of the crosslinker was added to around 100 µg of secondary antibodies and incubated at ambient temperature for 2 hours with mild shaking. Excess crosslinkers were removed by buffer exchange using the same centrifugal concentrators till the crosslinker concentration becomes at least 100 folds lower that the antibody concentration. Conjugation ratio of the crosslinker with antibody is tested using a NanoPhotometer® (Implen NP80). This reaction resulted in a conjugation ratio of around nine cross-linkers per Antibody.

Secondary antibody against anti-rabbit IgG () was conjugated to Cy5 similarly as described above. The antibody was buffer exchanged to 1× PBS and concentrated to over 1.5 mg/ml using centrifugal concentrators as mentioned above. Spins were performed at maximum recommended protocol of 12000 × g. Cy5-NHS Ester crosslinker (Jena Biosciences #CLK-A134-10) stock solution was stored at a concentration of 15.39 mM. A 25-fold excess of the crosslinker was added to around 100 µg of secondary antibodies and incubated at ambient temperature for 2 hours with mild shaking. Excess crosslinkers were removed by buffer exchange using the same centrifugal concentrators till the crosslinker concentration becomes at least 100 folds lower that the antibody concentration.

A 15× excess docking strand with azide-modification was added (Supplementary Table 1) to the cross-linker conjugated antibodies and incubated overnight at 4 °C under mild shaking. Excess docking strands was then removed using the centrifugal filters as mentioned above. The antibodies were stored at 4 °C until further use.

Nanobody Purification and Conjugation

Nanobodies were purified and conjugated as stated earlier.^3^ Plasmid (pTP1183) containing Anti-Rabbit IgG Fc nanobody sequence with a single cysteine (AddGene #104163) originally deposited by Dirk Görlich Lab was purchased. Plasmid were isolated from *E. coli* colonies grown on Luria Bertani Agar and in Luria Bertani Broth supplemented with 50 µg/ml of Kanamycin (Sigma #60615) using Qiagen Plasmid Extraction Mini-Prep Kit (Qiagen # 27106) using manufacturer’s protocol. The plasmids were then used to transform *E. coli* BL21 cells and grown on Luria Bertani Agar with 50 µg/ml of Kanamycin. Single colonies were picked for inoculating 25 ml of 2×YT (HiMedia #M1251-500G) primary culture with 50 µg/ml of Kanamycin at 28 °C overnight at 180 rpm. This culture was then added to 250 ml of 2×YT secondary culture at 25 °C at 180 rpm. The culture was then induced after one hour with 0.2 mM IPTG and incubated at 25 °C at 180 rpm for four and a half hours. Cells were then harvested on a Beckman Coulter centrifuge with the JA10 rotor at 8000 rpm at 4 °C. The pellets were stored at -80 °C.

The pellets were resuspended in lysis buffer and incubated on ice for 30 minutes prior to 1 hour sonication. The lysate was centrifuged at 30000 rpm on the above-mentioned equipment for 40 minutes at 4 °C. The supernatant was passed through a gravity-based Ni-NTA Affinity Purification Column (GBioscience 786-940). The column was washed thrice with an increasing concentration of imidazole in the lysis buffer (20 mM, 40 mM, and 80 mM). The protein was then eluted with elution buffer.

Buffer was exchanged to maleimide labelling buffer using Cytiva HiPrep 26/10 Desalting Column (#71-5004-28-EG) on a Bio-Rad NGC System. For tag cleavage, bdNEDP1 protease was added to the desalted eluant at a final concentration of 300 nM and incubated overnight at 4 °C. The tag and the protease were separated from the cleaved nanobodies using a gravity-based Ni-NTA affinity chromatography as stated earlier. The tag and the protease bind to the column and the nanobody is collected in the flow through. The single cysteines are then reduced using 10 mM DTT and 1 mM EDTA overnight at 4 °C. Desalting was repeated as stated above to remove the reducing agent that would hinder the reaction of crosslinker attachment.

Here we describe nanobody conjugation methodology.^4^ DBCO-PEG4-Maleimide (Jena Bioscience #CLK-A108P-10) crosslinker in DMSO was stored at a concentration of 59.2 mM. A five-fold molar excess of the crosslinker was added to the nanobodies an incubated overnight at 4 °C. Excess crosslinker was then removed using centrifugal filters (Merck Millipore Amicon Filter #UFC501096) using the manufacturer’s protocol. A three-fold molar excess of DNA with azide modification (Supplementary Table 1) was added to 100 µg of the nanobody carrying the crosslinker and incubated overnight at 4 °C for attaching DNA to the nanobodies.

Separation of labelled nanobodies from unlabeled nanobodies is essential to maximize labelling efficiency of the antibodies with nanobodies carrying docking strands. This separation was done using anion exchange column Resource Q (Cytiva #17117701). The column was equilibrated using low salt buffer. The nanobody-DNA mix was diluted using 0.5 × PBS to a volume of 250 µl and then injected onto the column. This was followed by a 5 ml wash with low salt buffer. Unlabeled nanobodies don’t bind to the column and come out in the flow through or washes. Low flow rate of 1 ml/minute was used with an increasing salt gradient of up to 50% high salt buffer over 20 ml. The DNA labelled nanobody elutes first followed by the free DNA molecules. The fraction containing labelled nanobody is then buffer exchanged to 1× PBS using Amicon centrifugal filters and aliquoted. The nanobodies are either flash frozen in liquid N_2_ and stored at -80 °C for long-term storage or stored at 4 °C for up to 1 month.

Cell Culture and Staining

HeLa Kyoto cells were cultured in Dulbecco's Modified Eagle Medium (DMEM Gibco #11995081) supplemented with 10% Fetal Bovine Serum (FBS Gibco #10270106) and 100 U/ml of Penicillin and 100 µg/ml of Streptomycin (Gibco #15140122) at 37 °C with 5% CO_2_. Cells were split when they reached ~70% confluency. For DNA-PAINT imaging experiments, cells were trypsinized with 0.05% Trypsin-EDTA (Gibco #25300054) for five minutes at 37 °C. Cells were then resuspended in antibiotic-free DMEM, centrifuged at 300× g at room temperature for 4 minutes and supernatant was discarded. The pellet was then resuspended in 1 ml of antibiotic-free DMEM. Cells were counted on a Countess™ 3 Automated Cell Counter (Thermofischer) and seeded on 8-well cover glass bottom dishes (CellVis #C8SB-1.5H) at a density of 8000 cells per well in antibiotic free DMEM. Cells were grown overnight as mentioned above.

For transcription inhibition experiments, Actinomycin D (ActD, Sigma-Aldrich #A9415) was dissolved to a final concentration of 4 mM in DMSO. After 12-16 hours of seeding, media in the wells was replaced with fresh media and incubated for thirty minutes. Prior to treatment, ActD was diluted to 4 μM in antibiotic-free DMEM supplemented with 10% FBS. DMSO was used as vehicle control. Then, media in the wells was replaced with ActD or DMSO containing media and incubated for 5 hours at 37 °C with 5% CO_2_. After this, the wells were washed thrice with 1 x PBS at 37 °C and immediately fixed with PFA fixation buffer at 37 °C.

For imaging single nuclear targets, media was removed from the well after 12 to 16 hours and the cells were fixed with prewarmed PFA fixation buffer for 10 minutes at 37 °C. Cells were then rinsed thrice with 1× PBS and stored at 4 °C until further use.

Chemically cross-linked cells were blocked and permeabilized using blocking buffer for 1 hour. This was followed by primary antibody staining for the respective targets at mentioned dilutions (Supplementary Table 1) and incubated for 4 hours at ambient temperatures. Sample were rinsed thrice with 1× PBS. Secondary nanobody carrying a single docking strand was used in separate wells for staining the samples. The samples were stained with a 1:200 dilution of secondary nanobodies in antibody incubation buffer for 45 minutes at ambient temperature. The sample was rinsed thrice with 1× PBS and was ready for DNA-PAINT imaging.

For 9-plex imaging of nuclear targets, primary antibodies (Supplementary Table 4) were pre-incubated with their respective docking strand conjugated secondary nanobodies in 10 μl of antibody incubation buffer at room temperature for 60 minutes. After pre-incubation, a 1:4 molar excess of unlabeled secondary nanobody was added to each mix and incubated for five minutes at room temperature. Following this, all eight pre-mixes were pooled together to a final volume of 100 μl. SC35 primary antibody (mouse-origin, Supplementary Table 4) was added to the mix and added to the permeabilized cells for one hour at room temperature. Next, the sample was washed five times with 1xPBS followed by a 45-minute incubation with docking strand conjugated anti-mouse secondary antibody. The sample was washed five times with 1×PBS followed by post-fixation with PFA fixation buffer at room temperature for ten minutes. The sample was rinsed thrice with 1× PBS and by quenching in quenching buffer for seven minutes. The sample was then rinsed thrice with 1× PBS, once with Buffer C before performing DNA-PAINT imaging. Samples were incubated with Gold Nanoparticles (Sigma #753688-25ML) at a dilution of 1:10 in buffer C for 10 minutes and then washed thrice with 1× PBS prior to imaging.

For imaging docking-strand conjugated nanobody alone controls, 1x PBS was used instead of primary antibodies and all following steps were followed exactly as described above for 9-plex nuclear target imaging. Imaging was performed by keeping the imaging parameters identical to a replicate of the mock dataset for direct comparison (Supplementary table 2).

For imaging the nucleolus, primary antibody against Fibrillarin (CST #2683T) was added to mock cells in antibody incubation buffer for 1 hour. The sample was washed five times with 1× PBS followed by incubation with Cy5-conjugated secondary nanobody for 1 hour. DNA-PAINT imaging was performed first with R3 imager (described below) followed by epifluorescence imaging at 640 nm excitation wavelength using Nikon D-LEDI Fluorescence LED illumination system.

Microscopy and Imaging

All imaging was performed on a Nikon Ti2 eclipse microscope body mated to a motorized H-TIRF unit and a perfect focus system (PFS). Teledyne Photometrics PRIME BSI sCMOS camera or Hamamatsu Fusion BT sCMOS camera was used for capturing images. Illumination with the 561 nm laser or the 640 nm laser was achieved using the L6cc laser combiner from Oxxius Inc., France. Oil immersion high NA objective lens capable of Total Internal Reflection Fluorescence (TIRF) imaging from Nikon (Nikon #Apo SR HP TIRF 100×, 1.49 NA, oil immersion) was used in total internal reflection condition for DNA origami nanostructure imaging. For nuclear targets and nuclear lamin, Highly Laminated Optical Sheet illumination was used. The camera was set with 2×2 – pixel binning along with cropping to effectively achieve a 66.56 × 66.56 µm^2^ field of view. Each pixel size was 130 nm × 130 nm.

A 20 × PCD (Protocatechuate 3,4-dioxygenase; Sigma #P8279-25UN) was prepared in 50 mM KCl, 1mM EDTA, and 100 mM Tris-Cl, pH 8.0, and 50% glycerol to a concentration of 6 µM. The stock PCD was then divided into 10 µl aliquots and stored at -20 °C for future use.

A 40 × PCA (Protocatechuic Acid / 3,4-Dihydroxybenzoic acid; Sigma #37580-100G-F) was made by dissolving 154 mg of PCA in 8 ml of Milli-Q® and 3 M NaOH (SRL #96311) was added dropwise and stirred until PCD completely dissolved. Volume was made up to 10 ml and 10 µl aliquots were stored at -20 °C for future use.

A 100× Trolox (6-hydroxy-2,5,7,8-tetramethylchroman-2-carboxylic acid; Sigma #238813-1G) was prepared by dissolving 100 mg of Trolox in 430 µl of methanol, 345 µl of 1 M NaOH and 3.2 ml of Milli-Q®. The solution was stored as 20 µl aliquots at -20 °C for future use.

A 500mM solution of Cysteamine (Sigma-Aldrich #M9768-25G) was prepared immediately before imaging by dissolving 38.5 mg of Cysteamine in 500 µl of nuclease free water (SRL #96370) and stored at 4 °C for no longer than 1 hour.

For multiplexed origami nanostructure imaging was performed by Exchange-PAINT^5^. In brief, a 200 µl of imaging buffer was prepared in buffer I+ with a final concentration of 1× PCA, 1× PCD and 1× Trolox, along with imagers. After each imaging round, a 1 ml of buffer I+ was used to wash the flow cell to remove the previous imager, and the next imager was introduced. This process was performed sequentially with all twelve imager sequences. Imaging was performed with the 561 nm laser. Imaging parameters are described in Supplementary table 2.

For kinetics studies, 100 µl of imaging buffer was prepared as mentioned earlier. The imaging buffer was added into the flow cell and sealed prior to imaging. Imaging with the 640 nm laser was performed prior to 561 nm laser to minimize effect of photobleaching of Atto647N fluorophore by the higher energy light source. Imaging parameters are described in Supplementary table 2.

For comparing imager behavior, imaging was performed with R1 and R8 imagers in two separate microfluidic channels. Each origami sample was first imaged at lower laser power (37 W/cm^2^) followed by imaging at a higher laser power (250 W/cm^2^). Imaging parameters are described in Supplementary table 2.

For cellular imaging, 200 µl of imaging buffer was prepared in buffer C with a final concentration of 1× PCA, 1× PCD, 1× Trolox and 8 mM cysteamine, along with imagers. For multiplexed cellular imaging, Exchange-PAINT was performed with all 12 imager sequences (9 imagers for nuclear 9-plex imaging) sequentially. Washes between each round were performed using at least 600 µl of 1 × PBS thrice. Imaging was performed with the 561 nm laser. Imaging parameters are described in Supplementary table 2.

Image Reconstruction and Kinetic Analysis

Image acquisition was performed using Nikon’s NIS Elements. Images were stored as *.nd2 files and read using Picasso Localize^2^ to obtain localizations. Drift correction was performed using redundant cross-correlation (RCC) followed by drift correction using picked origami nanostructures or fiducial markers using Picasso Render^2^. In multiplexed imaging, multiple channels were aligned to each other using RCC followed by picked fiducial markers.

Imager Crosstalk Analysis

All origamis were picked from the entire field of view and analyzed for the number of localizations coming from the same region during all imaging rounds. Random picks were generated at locations not closer than 260 nm from any other picks. Strip plots depicting the number of localizations arising from each channel over each origami were plotted to understand any obvious cross-talks (Supplementary Figure 3). Average localization counts greater than 10% of the correct target counts was considered a cross talk if it arises from an orthogonal target. A heatmap for the same was plotted (Figure 1e). The average values are tabulated in Supplementary Table 3. This analysis and plotting was performed using custom python codes.

Kinetic Analysis

Kinetic analysis was performed using inbuilt functions in Picasso Render^2^ where origamis showing signal for kinetic studies are picked individually and the properties of the picks are saved. The obtained files are then read using custom python codes and plotted. In brief, subsequent binding frames are clubbed together with a tolerance of 1 dark frame between two bright frames to define binding events. Length of each binding event is termed the bright times and the duration between binding events is termed the dark times.

Line Profile from Localization data

For line density plots of nuclear Lamin, localizations from rectangular regions were extracted and a frequency density plot was made along the length of the picked region. This was performed using custom python codes.

Resolution measurement for DNA origami structures

Origamis were picked and averaged using Picasso Average3 module. The localizations along the x-axis were then fit using 4 gaussians and along y-axis were fit with 3 gaussians. The average of the standard deviation of all the gaussian fits were computed to provide localization precision for each origami sets. This was performed using custom python codes.

Sampling measurement of DNA origami targets

Number of localizations on every origami at each docking strand was counted across frames. A complete target sampling was considered only when more than 20 localizations are present on the docking site. Target sampling was measured over time and was plotted. Heatmaps for 20 exemplary origamis representing the sampling evolution over every 1000 frames were plotted. This was performed using custom python codes.

Nuclear Data Extraction

Data from each cell was extracted from Picasso’s Render by picking a rectangular region encompassing the complete nucleus. Data was first cleaned for noise by eliminating localizations originating from binding events longer than 50 frames. The localizations from the Lamin B1 channel were then used to build a nuclear mask using AlphaShape library in Python after removing localizations from outside the nuclear region. The defined boundary was then used to filter all the other data channels to extract the nuclear localizations only.

Global Pearson’s Correlation

Nucleus was divided into sectors of size 130×130 nm (1 pixel size). Number of localizations from each sector for all channels were tabulated and Pearson’s correlation was calculated using custom Python code.

Spatial Correlation Viewpoint Analysis

Spatial correlation for each target localization with others was done as performed earlier^6^, concentric donuts of radii 100 nm to 1000 nm with a step size of 100 nm was used to count the number of localizations within each donut for both the species in question. Spearman’s correlation (𝜌) or Degree of Colocalization (DoC) for the number of localizations across 10 concentric donuts were then computed and assigned for each localization for the corresponding target in question.

These DoC values were then plotted as KDE plots for each target pair to express the trend of DoC values showcased by each target with the other. For viewpoint observations in Figure 4c, localizations and DoC values for the given region, highlighted by shaded area in the KDE plots, were extracted out and plotted as scatter plots. These are done using custom Python codes.

Pairwise cross-correlation analysis

Pairwise cross-correlation is a measure of the probability to find a target point in question at a given distance ‘d’, from any given localization of the viewpoint species. To calculate this, the distance between each point in the viewpoint species and all the target species are calculated and binned to 10 nm starting from d = 10 nm to d = 1000 nm. These counts are then normalized to the probability of finding the target point from a viewpoint if the targets are homogenously distributed within the same area. Post normalization, the values are plotted as a line plot from each viewpoint to every other target. These are done using custom Python codes.
