## Supplementary Figures for "High-Speed Multiplexed DNA-PAINT Imaging of Nuclear Organization using an Expanded Sequence Repertoire"

Supplementary Figure 1

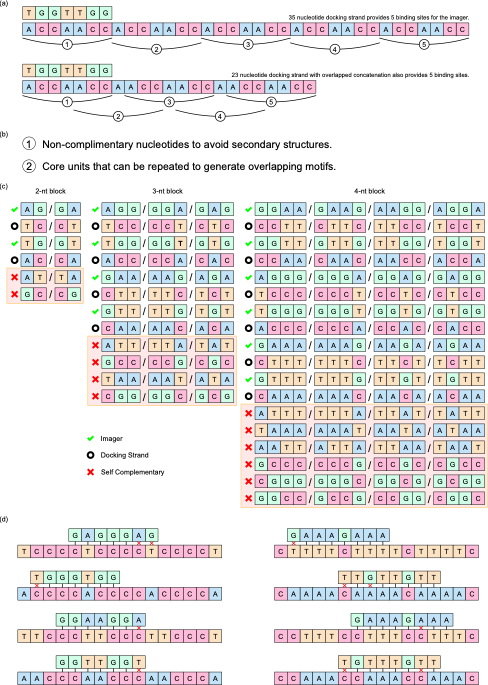

**Supplementary Figure 1: Fundamentals of good docking strand design and the usable sequence space.** (a) Concatenation in an overlapping manner can reduce the length of the docking strands without compromising on the available number of binding sites for the imager. (b) Rules that need to be satisfied for speed sequence building. (c) All possible two-nucleotide, three-nucleotide and four-nucleotide blocks possible. Green ticks indicate the sequences blocks used for imagers and the black circles indicate the sequences used for the docking strands. Red crosses indicate sequences that have self-complementarity and thus cannot be used. (d) Penta-nucleotide repeats show complementarity with other imagers with either one or two-nucleotide mismatches.

Supplementary Figure 2

**Supplementary Figure 2: 12-color DNA origami nanostructure imaging.** The figure showcases 50 unique DNA origami nanostructures from the same imaging field using different speed sequence imagers. (Scale Bar: 500 nm)

Supplementary Figure 3

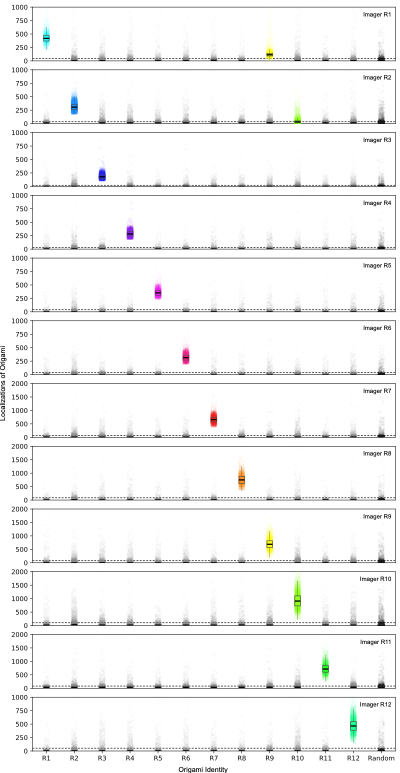

**Supplementary Figure 3: Number of localizations on each target origami.** Strip plot showing the number of localization on each DNA origami species with each imager. Dotted line shows 10% of the average number of localization on the correct target species (threshold). Coloured plots indicate that the mean is higher than the threshold for that imager-target combination.

Supplementary Figure 4

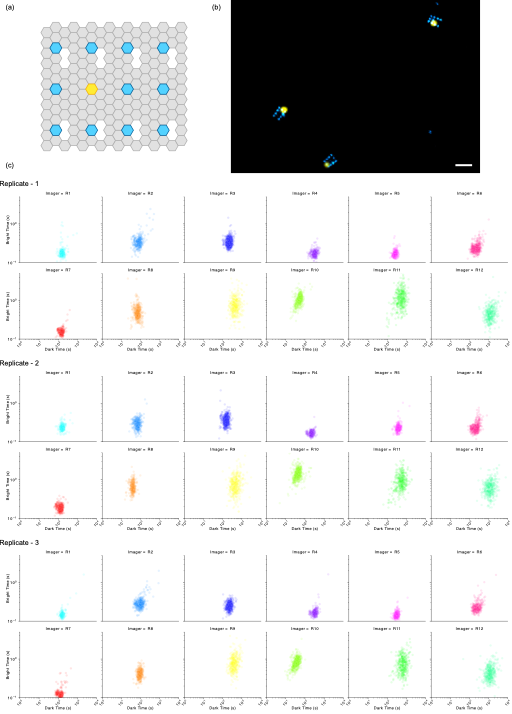

**Supplementary Figure 4: Kinetic analysis of speed sequence imagers.** (a) Graphical representation of the DNA origami nanostructure design of kinetic analysis. Blue hexagons represent the docking strands used for detecting the location of the DNA origami nanostructures. The yellow hexagon represents the site of the docking strand where kinetic analysis is being performed. Two colour DNA-PAINT imaging was used to image both the sets of docking strands sequentially. (b) Representative merged field of kinetic analysis. (Scale Bar: 100 nm) (c) Scatter plots representing the bright and dark times of all different imagers per docking strand per nanomolar concentration of imager.

Supplementary Figure 5

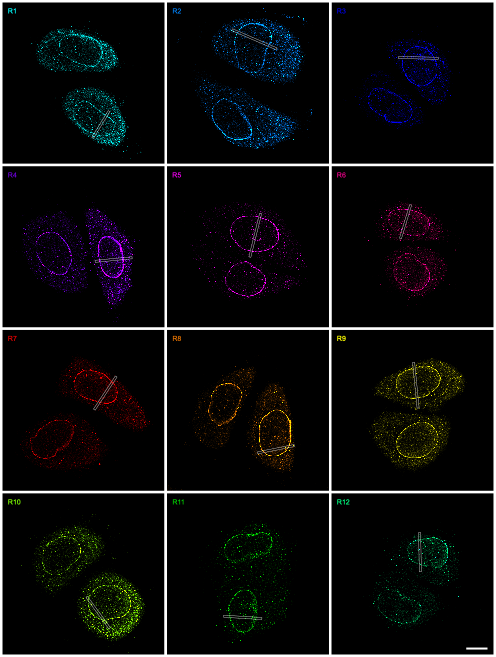

**Supplementary Figure 5: Nuclear Lamin staining with all 12 speed sequences.** Staining was done with primary antibody and secondary nanobody. Line profile plots for the marked regions are shown in Supplementary Figure 6. (Scale Bar: 10 µm)

Supplementary Figure 6

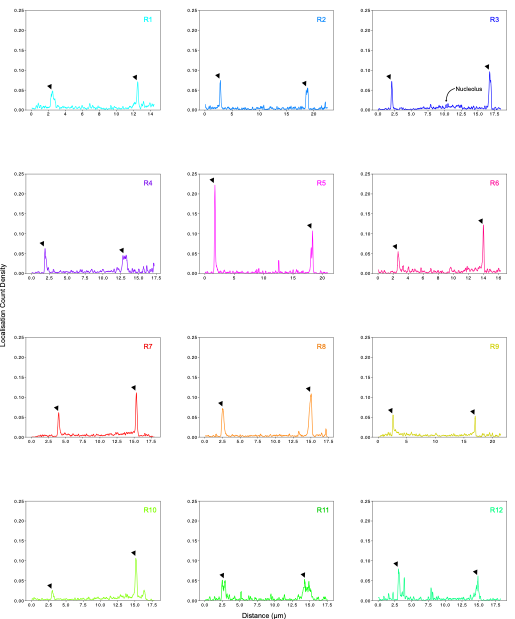

**Supplementary Figure 6: Line profile plots of Nuclear Lamin regions.** Plots are drawn along the regions marked in Supplementary Figure 5. Black arrows point to the peaks arising from LaminB1 signal. Imager R3 shows increased intra-nuclear signal likely due to non-specific signals from the nucleolus.

Supplementary Figure 7

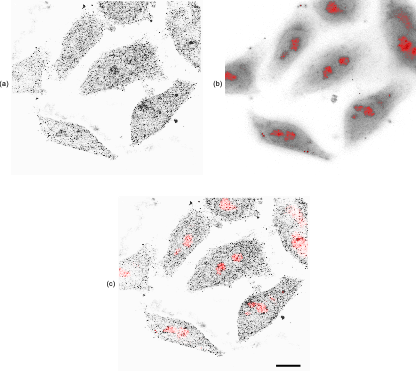

**Supplementary Figure 7: R3 imager shows off-target binding to the nucleolus. (**a) DNA-PAINT image with R3 imager alone (10000 frames; 1000 s); (b) Immunofluorescence image of Fibrillarin using Cy5 labelled secondary antibody; (c) Overlay of (a) and (b); (Scale bar: 10 µm)

Supplementary Figure 8

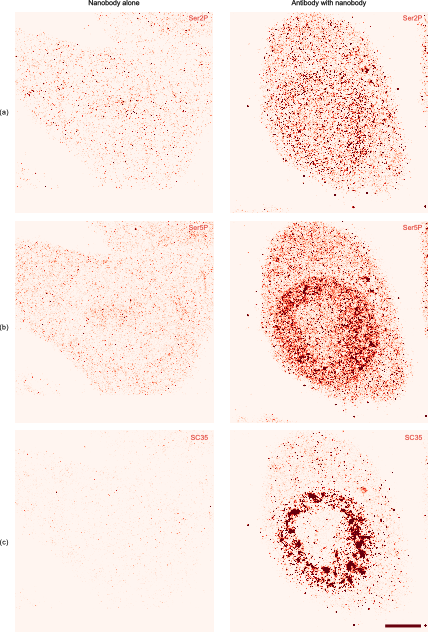

**Supplementary Figure 8: Nanobody-docking strand alone controls show negligible off-target imager localizations in the nucleus.** DNA-PAINT images Left: Secondary nanobody alone; Right: Corresponding primary antibody-secondary nanobody complex. Targets: (a) RNAPolII-S2P-R7 imager, (b) RNAPolII-S5P-R1 imager, (c) SC35-R8 imager. Imaging parameters were identical for each pair. (Scale bar: 5 µm)

Supplementary Figure 9

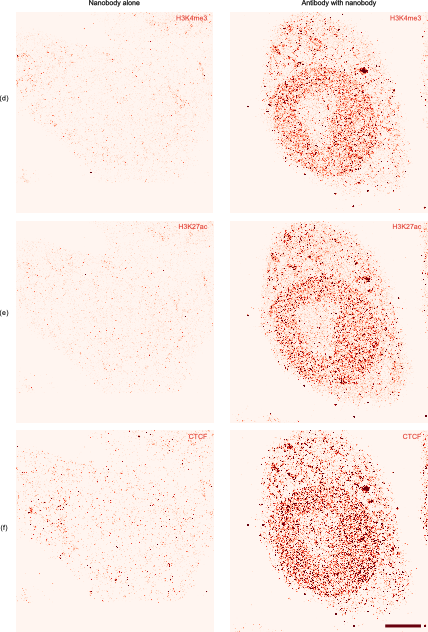

**Supplementary Figure 9: Nanobody-docking strand alone controls show negligible off-target imager localizations in the nucleus.** DNA-PAINT images Left: Secondary nanobody alone; Right: Corresponding primary antibody-secondary nanobody complex. Targets: (d) H3K4me3-R4 imager, (b) H3K27ac-R2 imager, (d) CTCF-R5 imager. (Scale bar: 5 µm)

Supplementary Figure 10

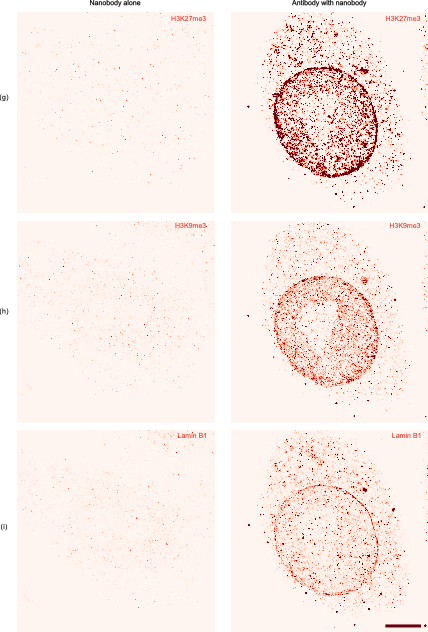

**Supplementary Figure 10: Nanobody-docking strand alone controls show negligible off-target imager localizations in the nucleus.** DNA-PAINT images Left: Secondary nanobody alone; Right: Corresponding primary antibody-secondary nanobody complex. Targets: (g) H3K27me3-R11 imager, (h) H3K9me3-R12 imager, (i) LaminB1-R6 imager. (Scale bar: 5 µm)

Supplementary Figure 11

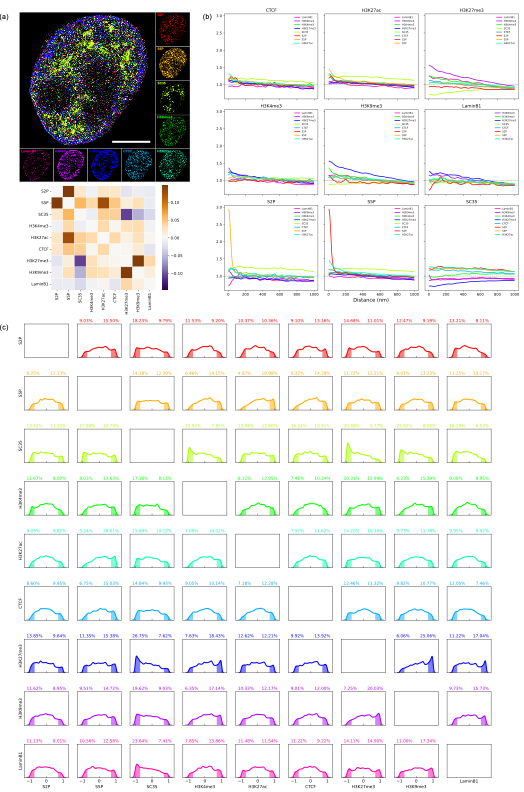

**Supplementary Figure 11: 9-plex imaging and analysis of Mock-treated Hela Kyoto cells**. (a) Top: 9-plex DNA-PAINT Image, Bottom: Global Pearson’s correlation analysis for all 9 nuclear targets; (b) PCCF analysis with all 9 targets as the viewpoints; (c) KDE plots representing DoC analysis of each target with respect to the other 8 targets. (Scale bar: 5 µm)

Supplementary Figure 12

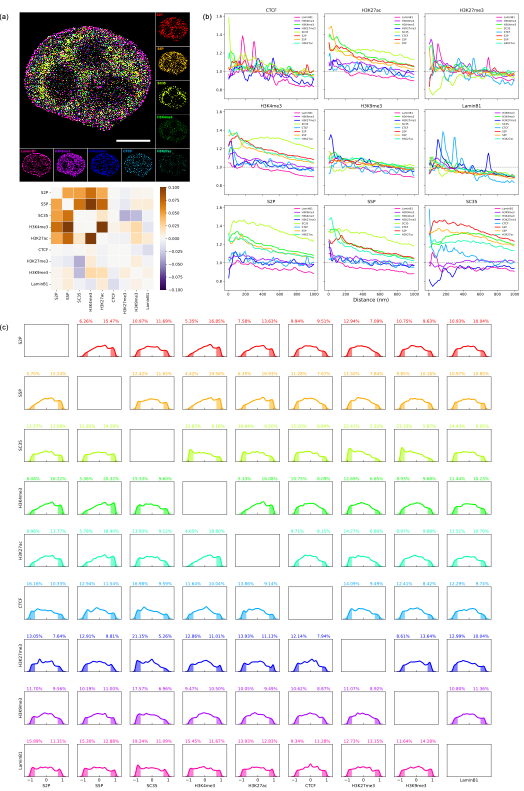

**Supplementary Figure 12: 9-plex imaging and analysis of Mock-treated Hela Kyoto cells**. (a) Top: 9-plex DNA-PAINT Image, Bottom: Global Pearson’s correlation analysis for all 9 nuclear targets; (b) PCCF analysis with all 9 targets as the viewpoints; (c) KDE plots representing DoC analysis of each target with respect to the other 8 targets. (Scale bar: 5 µm)

Supplementary Figure 13

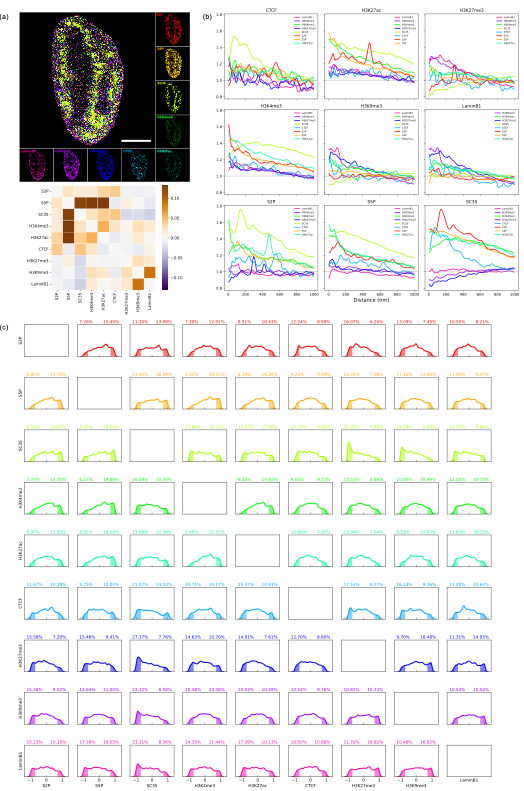

**Supplementary Figure 13: 9-plex imaging and analysis of Mock-treated Hela Kyoto cells**. (a) Top: 9-plex DNA-PAINT Image, Bottom: Global Pearson’s correlation analysis for all 9 nuclear targets; (b) PCCF analysis with all 9 targets as the viewpoints; (c) KDE plots representing DoC analysis of each target with respect to the other 8 targets. (Scale bar: 5 µm)

Supplementary Figure 14

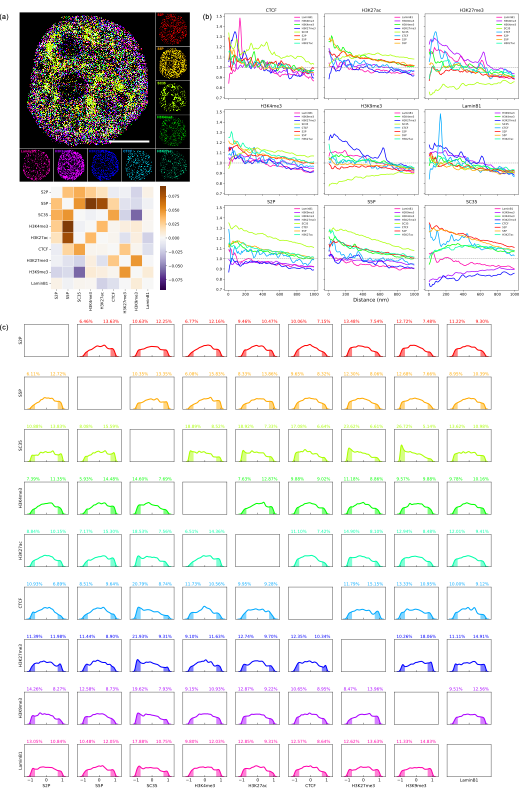

**Supplementary Figure 14: 9-plex imaging and analysis of Mock-treated Hela Kyoto cells**. (a) Top: 9-plex DNA-PAINT Image, Bottom: Global Pearson’s correlation analysis for all 9 nuclear targets; (b) PCCF analysis with all 9 targets as the viewpoints; (c) KDE plots representing DoC analysis of each target with respect to the other 8 targets. (Scale bar: 5 µm)

Supplementary Figure 15

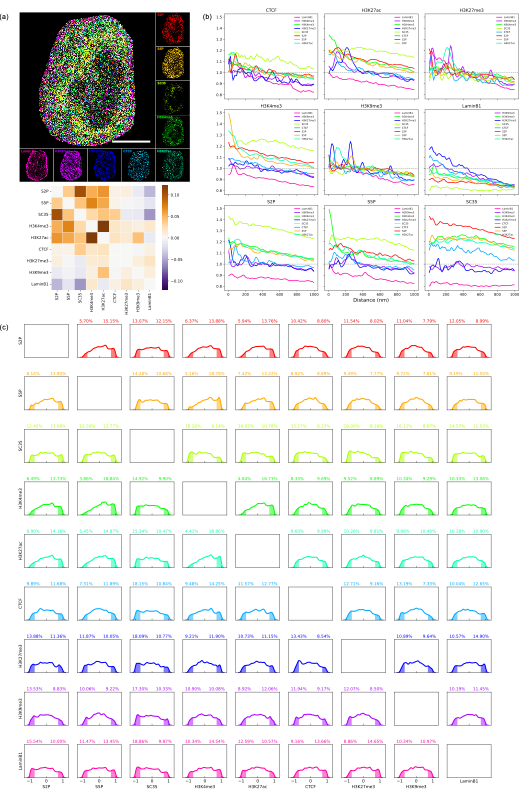

**Supplementary Figure 15: 9-plex imaging and analysis of Mock-treated Hela Kyoto cells**. (a) Top: 9-plex DNA-PAINT Image, Bottom: Global Pearson’s correlation analysis for all 9 nuclear targets; (b) PCCF analysis with all 9 targets as the viewpoints; (c) KDE plots representing DoC analysis of each target with respect to the other 8 targets. (Scale bar: 5 µm)

Supplementary Figure 16

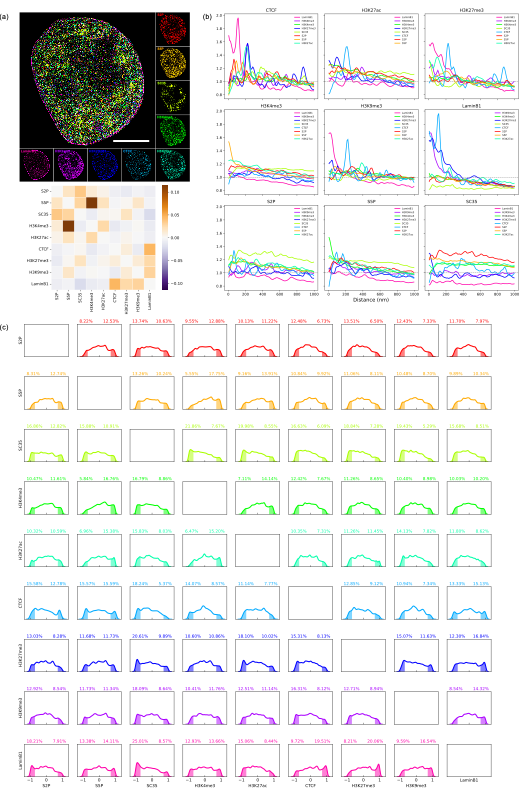

**Supplementary Figure 16: 9-plex imaging and analysis of Mock-treated Hela Kyoto cells**. (a) Top: 9-plex DNA-PAINT Image, Bottom: Global Pearson’s correlation analysis for all 9 nuclear targets; (b) PCCF analysis with all 9 targets as the viewpoints; (c) KDE plots representing DoC analysis of each target with respect to the other 8 targets. (Scale bar: 5 µm)

Supplementary Figure 17

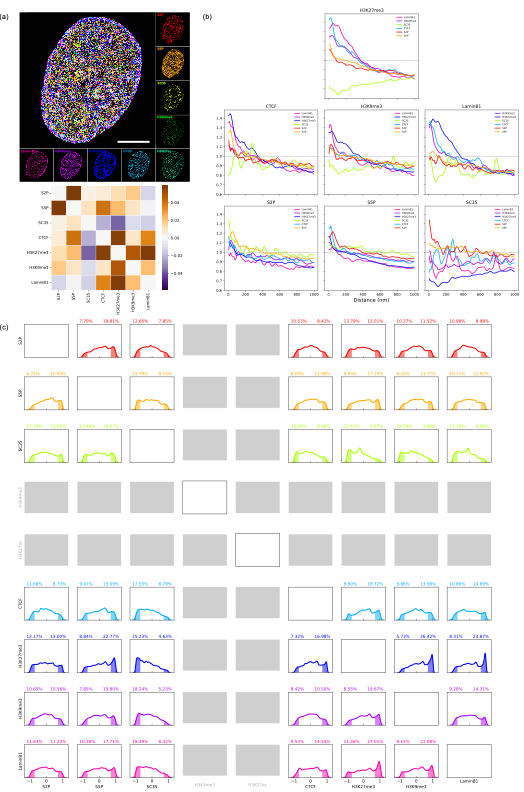

**Supplementary Figure 17: 9-plex imaging and analysis of ActD-treated Hela Kyoto cells**. (a) Top: 9-plex DNA-PAINT Image, Bottom: Global Pearson’s correlation analysis for all 9 nuclear targets; (b) PCCF analysis with all 9 targets as the viewpoints; (c) KDE plots representing DoC analysis of each target with respect to the other 8 targets. (Scale bar: 5 µm)

Supplementary Figure 18

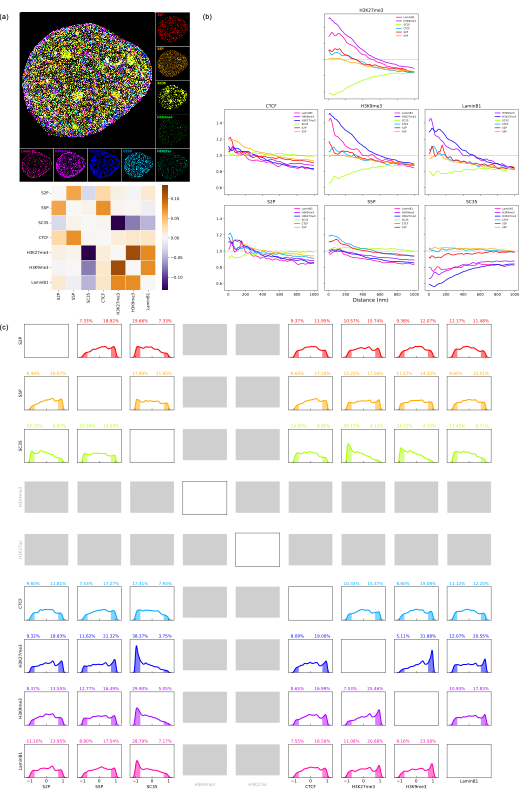

**Supplementary Figure 18: 9-plex imaging and analysis of ActD-treated Hela Kyoto cells**. (a) Top: 9-plex DNA-PAINT Image, Bottom: Global Pearson’s correlation analysis for all 9 nuclear targets; (b) PCCF analysis with all 9 targets as the viewpoints; (c) KDE plots representing DoC analysis of each target with respect to the other 8 targets. (Scale bar: 5 µm)

Supplementary Figure 19

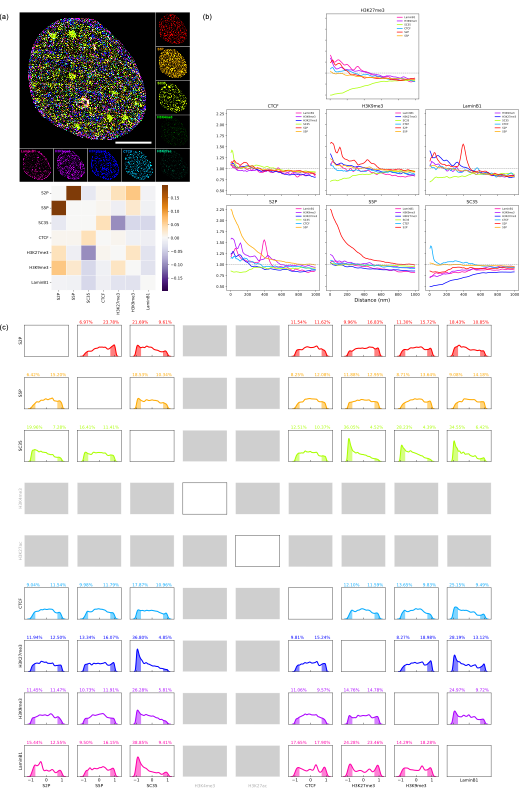

**Supplementary Figure 19: 9-plex imaging and analysis of ActD-treated Hela Kyoto cells**. (a) Top: 9-plex DNA-PAINT Image, Bottom: Global Pearson’s correlation analysis for all 9 nuclear targets; (b) PCCF analysis with all 9 targets as the viewpoints; (c) KDE plots representing DoC analysis of each target with respect to the other 8 targets. (Scale bar: 5 µm)

Supplementary Figure 20

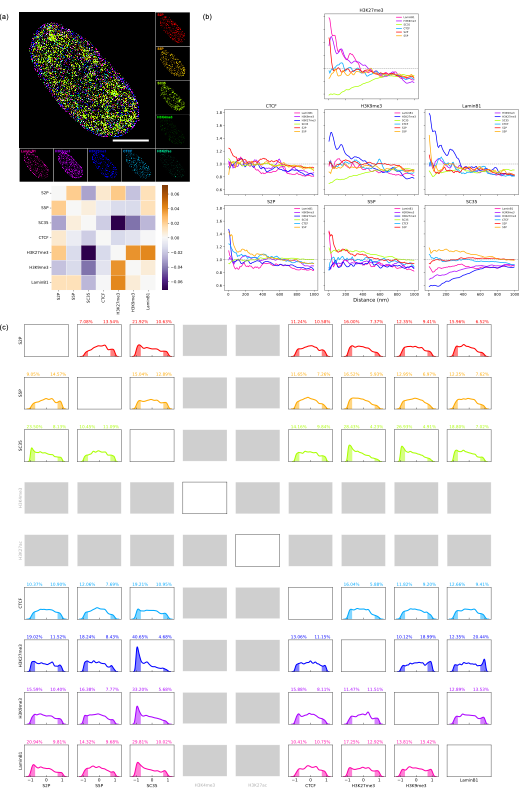

**Supplementary Figure 20: 9-plex imaging and analysis of ActD-treated Hela Kyoto cells**. (a) Top: 9-plex DNA-PAINT Image, Bottom: Global Pearson’s correlation analysis for all 9 nuclear targets; (b) PCCF analysis with all 9 targets as the viewpoints; (c) KDE plots representing DoC analysis of each target with respect to the other 8 targets. (Scale bar: 5 µm)

Supplementary Figure 21

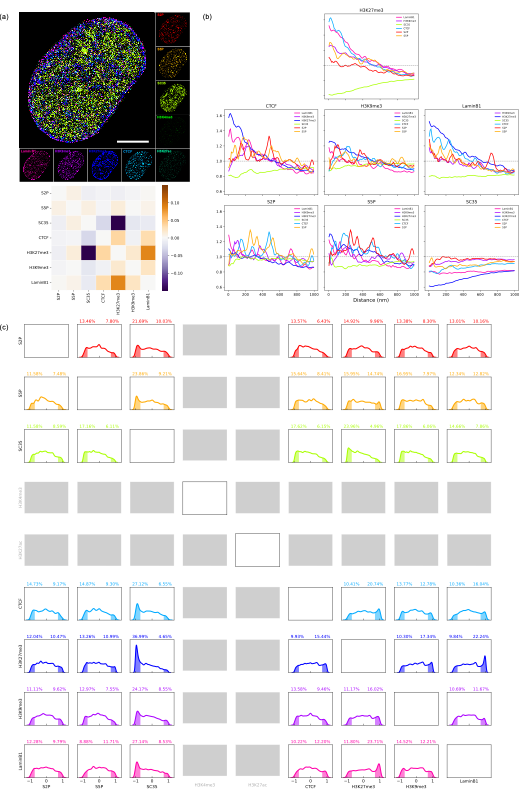

**Supplementary Figure 21: 9-plex imaging and analysis of ActD-treated Hela Kyoto cells**. (a) Top: 9-plex DNA-PAINT Image, Bottom: Global Pearson’s correlation analysis for all 9 nuclear targets; (b) PCCF analysis with all 9 targets as the viewpoints; (c) KDE plots representing DoC analysis of each target with respect to the other 8 targets. (Scale bar: 5 µm)

Supplementary Figure 22

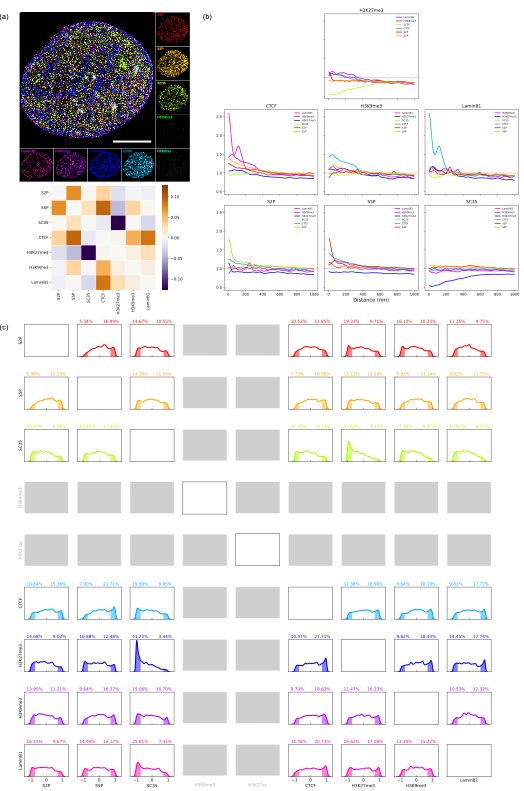

**Supplementary Figure 22: 9-plex imaging and analysis of ActD-treated Hela Kyoto cells**. (a) Top: 9-plex DNA-PAINT Image, Bottom: Global Pearson’s correlation analysis for all 9 nuclear targets; (b) PCCF analysis with all 9 targets as the viewpoints; (c) KDE plots representing DoC analysis of each target with respect to the other 8 targets. (Scale bar: 5 µm)

Supplementary Figure 23

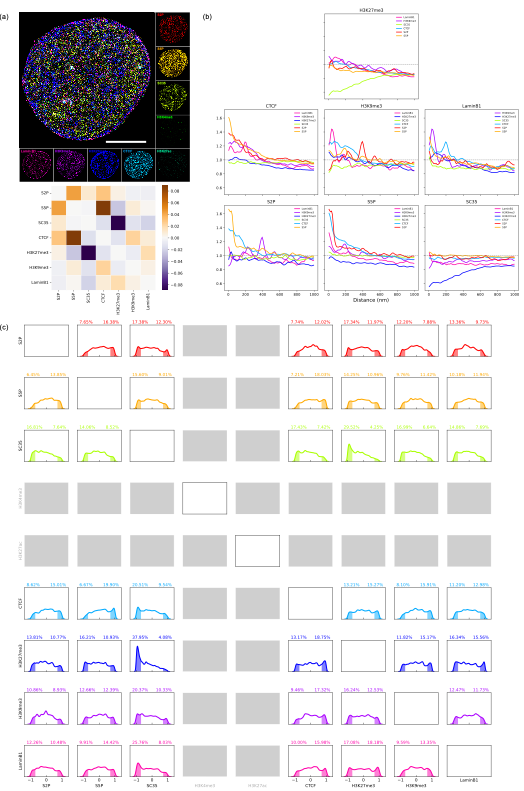

**Supplementary Figure 23: 9-plex imaging and analysis of ActD-treated Hela Kyoto cells**. (a) Top: 9-plex DNA-PAINT Image, Bottom: Global Pearson’s correlation analysis for all 9 nuclear targets; (b) PCCF analysis with all 9 targets as the viewpoints; (c) KDE plots representing DoC analysis of each target with respect to the other 8 targets. (Scale bar: 5 µm)

Supplementary Figure 24

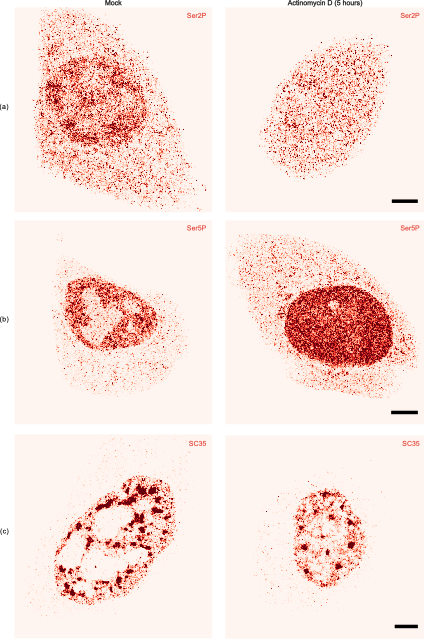

**Supplementary Figure 24: DNA-PAINT imaging of individual targets in Mock (left) and ActD (right) treated cells.** Imaging parameters were identical for each pair. (a) RNAPolII-S2P-R7, (b) RNAPolII-S5P-R1, (c) SC35-R8**.** (Scale bar: 5 µm)

Supplementary Figure 25

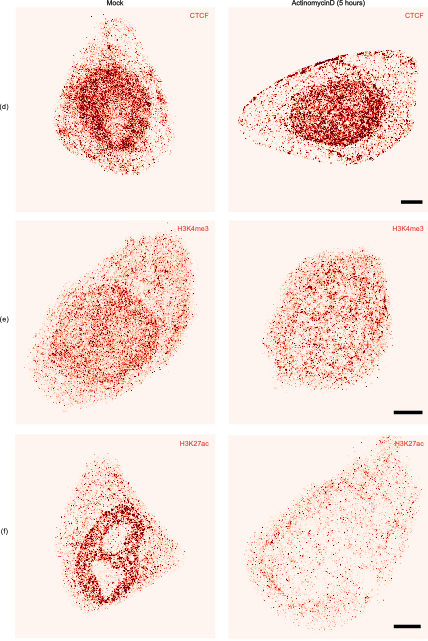

**Supplementary Figure 25: DNA-PAINT imaging of individual targets in Mock (left) and ActD (right) treated cells.** Imaging parameters were identical for each pair. (d) CTCF-R5, (e) H3K4me3-R4, (f) H3K27ac-R2**.** (Scale bar: 5 µm)

Supplementary Figure 26

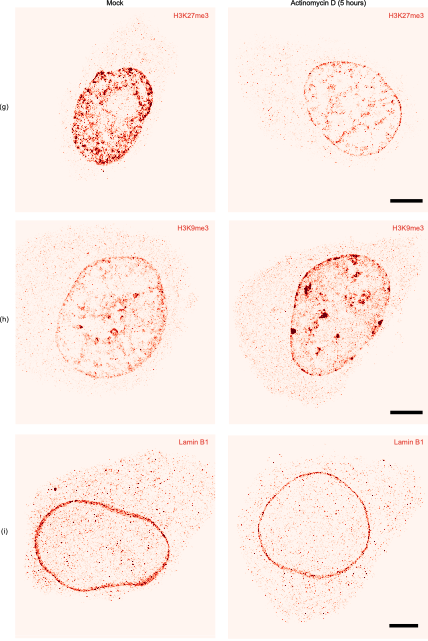

**Supplementary Figure 26: DNA-PAINT imaging of individual targets in Mock (left) and ActD (right) treated cells.** Imaging parameters were identical for each pair. (g) H3K27me3-R11, (h) H3K9me3-R12, (i) LaminB1-R6 (Scale bar: 5 µm)
